# Connecting myelin-related and synaptic dysfunction in schizophrenia with SNP-rich gene expression hubs

**DOI:** 10.1101/080044

**Authors:** Hedi Hegyi

## Abstract

The recent availability of several genome-wide data sets such as genome-wide mapping of SNP-rich regions and differentially methylated genes in schizophrenic individuals and gene expression data in all brain compartments across the span of human life prompted us to integrate these datasets to gain a better insight into the underlying gene networks driving this enigmatic disease.

We summed up the differentially methylated “expression neighbors” (i.e. genes with positively or negatively correlating expression values) of genes that fall into one of 108 distinct schizophrenia-associated genetic loci with high number of SNPs in schizophrenic patients derived from a large cohort of pooled sequencing experiments. Surprisingly, the number of expression neighbors (with a Pearson correlation of R>=0.8 or R<=−0.7) of the genes falling into the 108 genomic regions were about **35 times** higher for the positively correlating genes and **32 times** higher for the negatively correlating ones than for the rest of the ~16000 genes outside these loci. While the genes in the 108 loci have relatively little known impact in schizophrenia, using this approach we identified many more known schizophrenia-related important genes with a high degree of connectedness to other genes and high scores of differentially methylated probes for their expression neighbors (such as MBP, MOBP, GRIA1, COMT, SYNGR1, MAP2 and DGCR6), validating our approach.

The analysis revealed that the most positively correlating as well as the most negatively correlating genes affect synapse-related genes the most, offering an explanation and a unified view into the root cause of schizophrenia.

## Introduction

Gene expression correlation, protein-protein interaction and other high-throughput experiments in the post-genomic era have revealed that genes tend to form complex, scale-free networks where most genes have a few connections with others and a few have a high number of interactions, establishing them as important hubs in these gene networks [1]. These highly interconnected genes have become the targets of intense research expecting them to play prominent roles in genetic diseases. However, measurements found only a weak correlation between disease genes and hubs, e.g. Barabasi et al. [2] found that disease genes have 32% more interactions with other proteins than non-disease proteins, arguing that genetic mutations in topologically central, widely expressed genes are more likely to result in severe impairment of normal development, leading to lethality in utero and eventual deletion from the population [2].

This is apparently not the case in schizophrenia, a disease that steadily affects about one percent of the population despite the lower fecundity of the affected individuals [3]. In most cases of schizophrenia no known mutations exist in protein-coding genes [4] and the role of gene expression dysregulation in schizophrenia is increasingly recognized [5, 6].

A recent meta-analysis, examining the mutations in the genomes of 36,000 schizophrenics, identified 108 distinct genomic regions with significantly higher mutation rates [7]. We integrated this set with gene expression data from the Allen Brain Atlas. After determining the correlating and anti-correlating neighbors of genes in the 108 loci we found that the median number of correlating neighbors were about **35 times** higher for the genes in the 108 regions than in the rest of the genome (**32 times** for the anti-correlating pairs).

We also integrated the data with a recent methylome study in schizophrenics [8]. Ranking the genes for the hypermethylated probes of the positively and negatively correlating gene pairs identified the top gene as SYNGR1, a synapse-related gene whose regulatory region overlaps with one of the 108 SNP-rich loci in [7]. We also identified MBP and MOBP as two highly ranked genes in the anti-correlating set. They are both myelin-related, anti-correlating with a large number of synapse-related and glutamate receptor genes, offering a model that connects these frequently observed but so far disjoint pathologies in schizophrenia.

## Results

### Genes in the most mutated genomic regions have the highest number of correlating partners

In the first step we identified all human gene expression pairs with a Pearson correlation of >=0.8 or <=−0.7 in human brain tissues using the Allen Brain Atlas (website: brain-map.org), [9, 10] and a recent database with pre-calculated correlations for all relevant gene pairs (website: SZDB.org) [11]. This resulted in 1,257,407 positively correlating and 1,108,585 negatively correlating unique gene pairs belonging to 16829 and 16761 individual genes, respectively (**Supplementary File 1**).

Subsequently we mapped all known human genes in 108 highly mutated human genomic regions identified in [7] separately for genes whose promoters or only *cis* regulatory elements fall into these 108 loci. Filtering for genes present in the correlating or anti-correlating pairs in **Supplementary File 1** resulted in 254 promoter-selected and 462 cis-selected genes.

Counting the positively and negatively correlating partners for all the genes except in the 108 loci in [7] resulted in a median number of 71 positively and 63 negatively correlating pairs, respectively (Figure 1). Unexpectedly, the median numbers of correlating pairs for the promoter-selected genes were 2472 and 2013, corresponding to a **35**-fold and **32**-fold increase for the positive and negative pairs, respectively.

**Figure 1.**
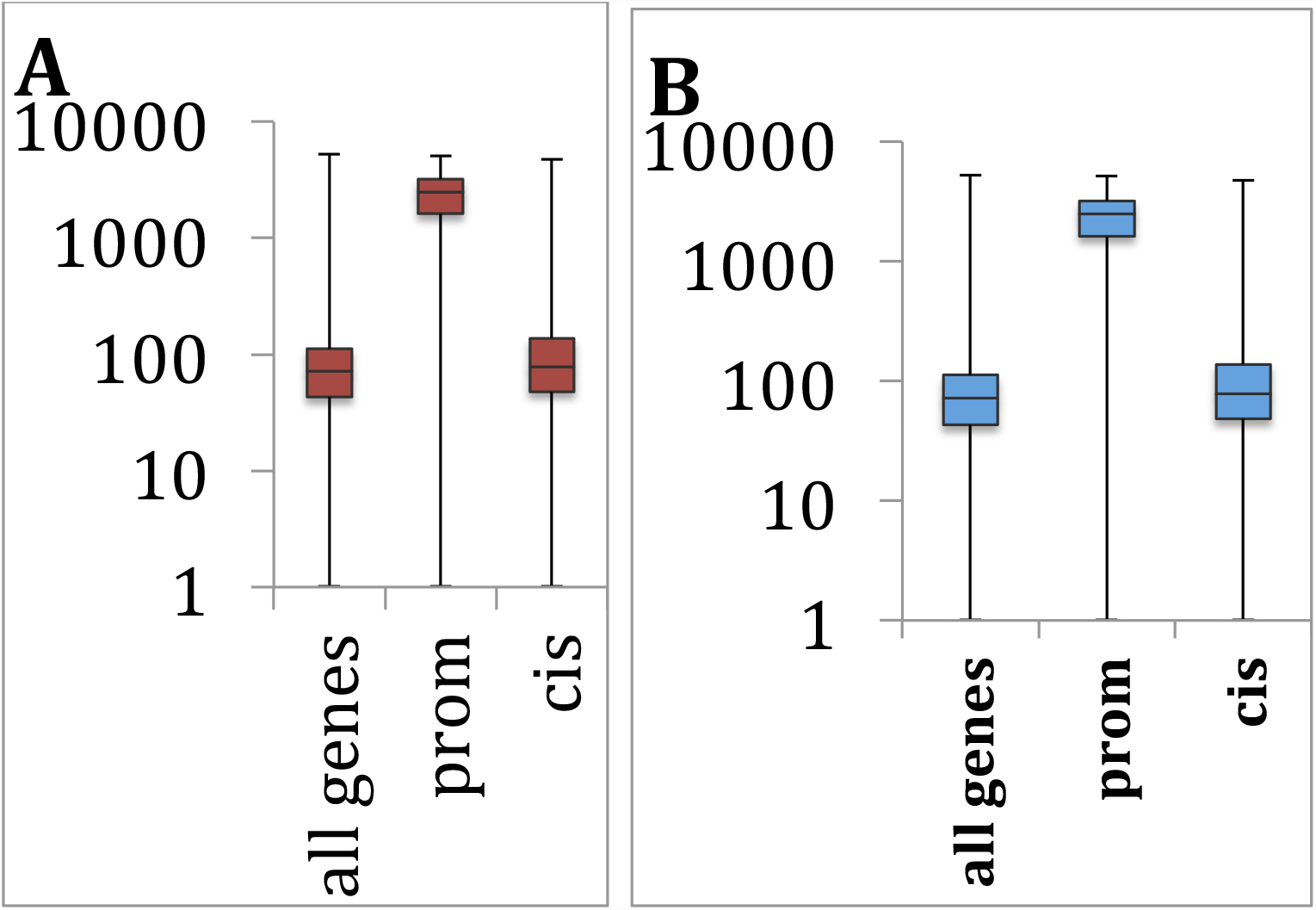
Boxplots of inteacting partners for all genes and promoter-selected and cis-selected genes in 108 SNP-rich genomic regions taken from [7]. (**A**) positive, (**B**) negative correlating partners.

The number of correlating pairs for the cis-selected genes also proved to be higher (386 and 320 pairs on average for positively and negatively correlating pairs, respectively), corresponding to an approximately 2.5-fold increase when compared to the general gene population. The median values were similar to the average gene population in this category (Figure 1) but an unpaired t-test showed that the *cis*-derived genes and the general gene population (excluding the 108-loci genes) are significantly different (p-value=0.012).

We also investigated the number of differentially methylated gene pairs using a table of 56001 differentially methylated probes from a genome-wide methylome study in schizophrenics [8]. We counted separately the number of hypermethylated and hypomethylated genes for the highly correlating and anti-correlating gene pairs and also the number of hyper- and hypomethylated probes belonging to these genes. The sums of scores for both the number of differentially methylated gene pairs and the total sum of their differentially methylated probes are shown in **Supplementary File 2** separately for the positively and negatively correlating pairs.

To determine which measurement distinguishes the best between schizophrenia-related and non-specific genes we calculated the effect size for the 108-loci genes and also for genes annotated by Genecards and Malacards [12, 13] as schizophrenia-related (Figure 2), using Cliff's delta for five different measurements (see legend for details). All the underlying data are shown in **Supplementary File 2**.

**Figure 2.**
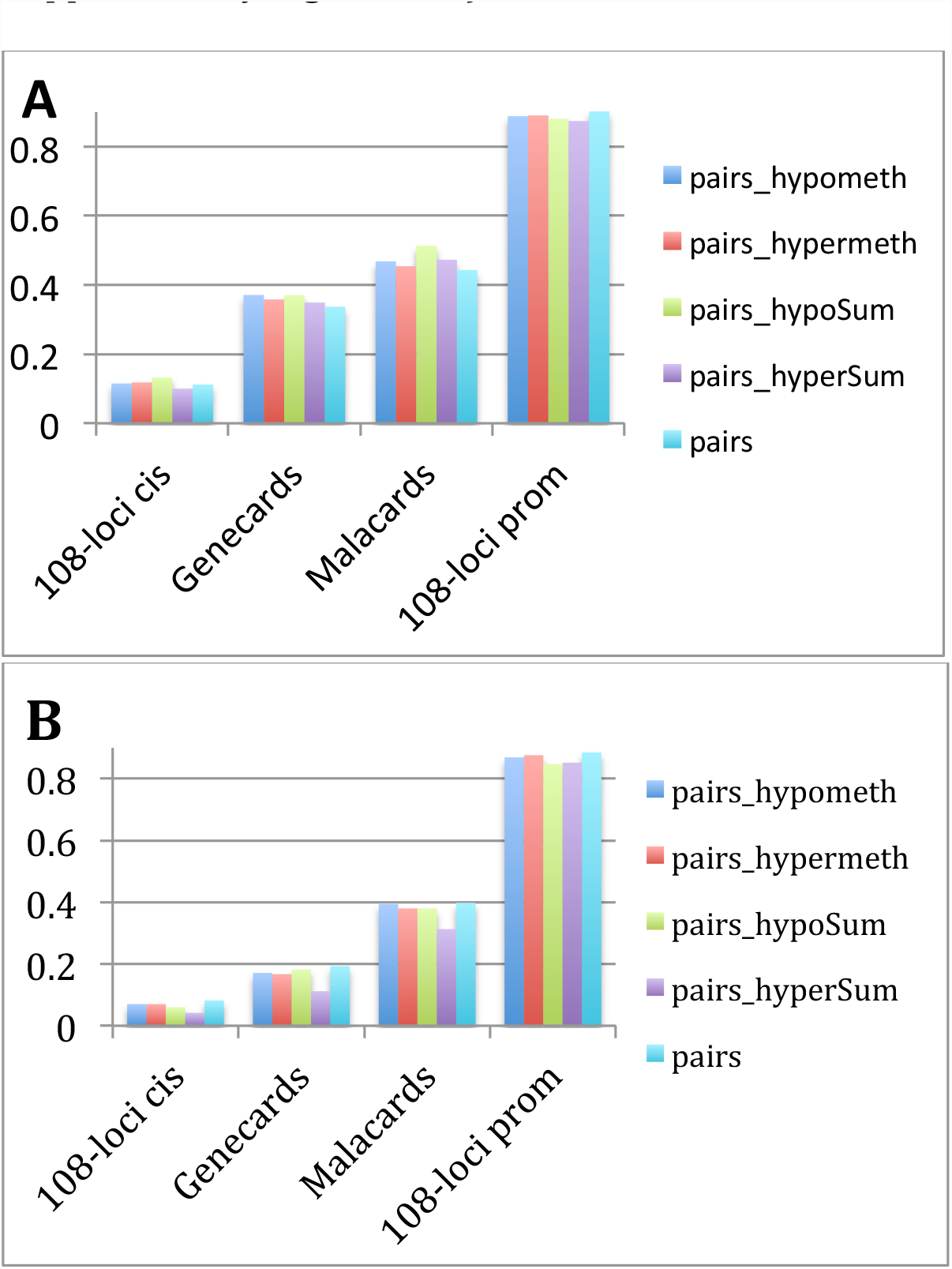
The **effect sizes** for the ratios of gene neighbor numbers between **specific** genes and their **complementary** gene sets (i.e. the rest of the ~16k genes present in the study): 108-loci cis genes; Malacards-annotated schizophrenia genes; Genecards-annotated schizophrenia genes; 108-loci promoter genes. Effect sizes were calculated for 5 different measurements: (i) **pairs**, the total number of correlating gene pairs; (ii) **pairs_hypometh**, the number of correlating gene pairs that are hypomethylated (defined as in at least one probe the gene is hypomethylated); (iii) **pairs_hypermeth**, the number of correlating gene pairs that are hypermethylated; (iv) **pairs_hypoSum**, the total number of hypomethylated probes of the correlating gene pairs; (v) **pairs_hyperSum**, the total number of hypermethylated pairs for the correlating gene pairs. The numbers were filtered using only pairs of genes in **Supplementary Table 1 A** & B where both genes were differentially methylated in [8]. (**A**) Positively, (**B**) Negatively correlating gene pairs.

Figure 2 a and b shows the effect sizes for all relevant pairs of complementary gene sets (i.e. promoter-derived *vs.* non-promoter-derived, Genecards *vs.* non-Genecards genes, etc.) comparing the numbers of correlating gene partners in one set to the numbers of the gene partners in the other gene set. As expected the promoter-derived 108-loci genes had the greatest effect size, followed by Malacards and Genecards, the 108-loci cis-derived genes having the smallest effect. The results were similar for the negatively correlating pairs (Figure 2b). Here we also filtered for gene pairs that were both differentially methylated in [8], which increased the effect size by 0.039 for “pairs” in Figure 2 for the promoter-derived positive set and by 0.061 for the same for the negative set. (Unfiltered charts shown in Supplementary Figure1 AB.)

### Specific genes with high numbers of correlating partners

We ranked all genes according to the number of hypermethylated probes (taken from [8]) of the correlating gene partners. Table 1 shows the top 20 genes with the most hypermethylated probes, in the positively and negatively correlating partners, respectively. The top-ranking gene in Table 1A is **SYNGR1**, synaptogyrin, a synapse-related gene, followed by **GRIN1**, a glutamate receptor and **CHRNB2**, a cholinergic receptor. All three genes have synapse-related functions, SYNGR1 and GRIN1 regulate synaptic plasticity whereas CHRNB2 regulates synapse assembly [14].

**Table 1.**
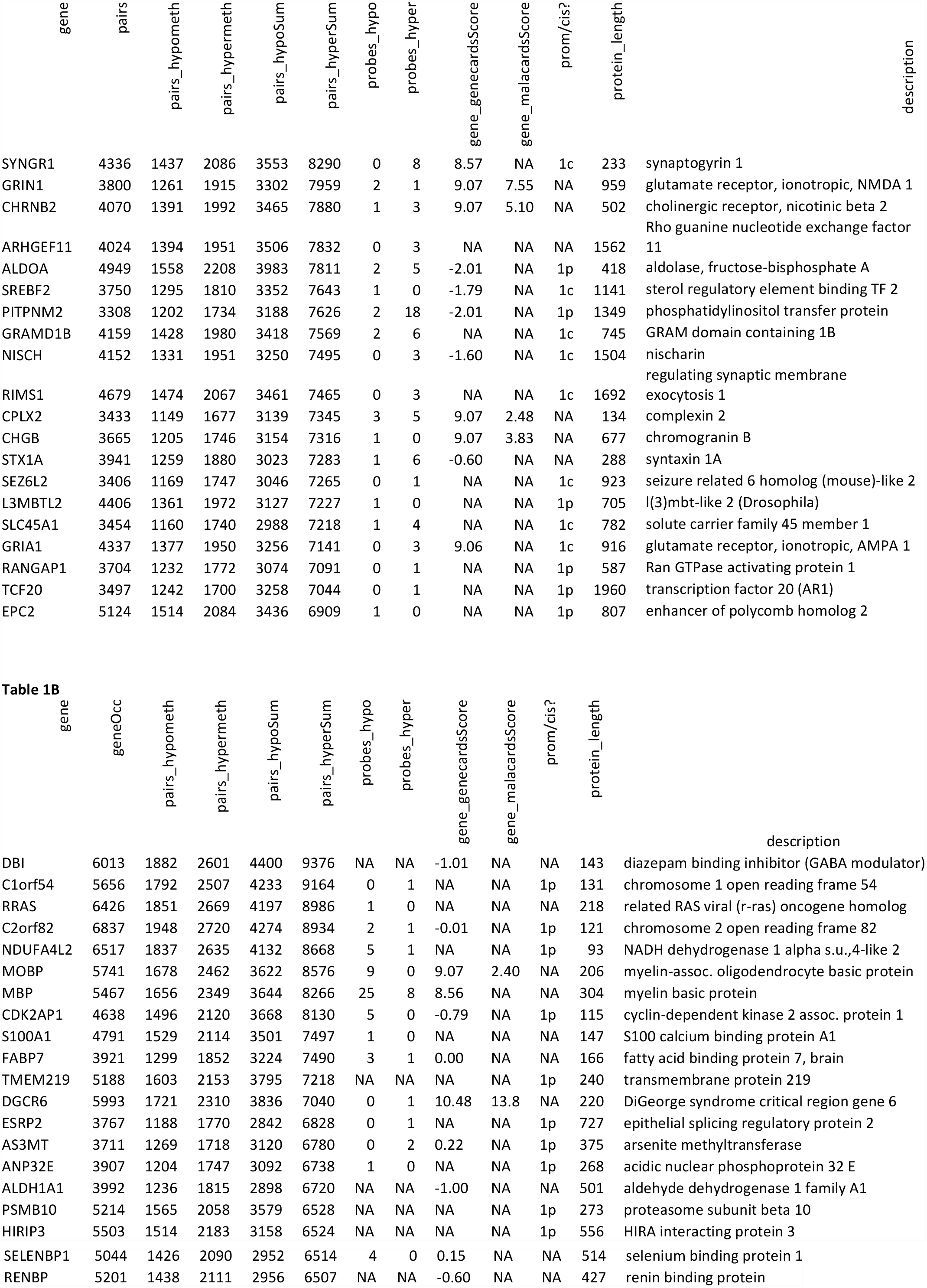
**(A)** The top 20 positively and **(B)** negatively correlating genes, ranked according to the hypermethylated probes (listed in column 6) in the correlating genes partners. The column “prom/cis?” indicates if the promoter of the gene (“1p”) or the cis regulatory region of the gene (“1c”) falls into the 108 highly mutated loci (see more details in the text).

SYNGR1 is also present in the 108 most mutated loci in [7] although only for its regulatory region, not for its promoter. Further functional analysis identified 5 more synapse-related genes in Table 1A: **NISCH** regulates synaptic transmission, **RIMS1** and **STX1A** regulate postsynaptic potential, **CPLX2** regulates synaptic plasticity and **GRIA1** regulates long-term synaptic depression.

Altogether 11 out of the 20 genes are annotated in Genecards as schizophrenia-related and 14 appear in the 108 genomic regions in [7]. The only gene in Table 1A that appears in neither is **ARHGEF11**, a glutamate transport enhancer, however it had a significantly higher expression in the thalamus of schizophrenics in [15].

**Table 1B** lists the top 20 genes with the highest number of hypermethylated probes for the negatively correlating partners. The top-ranking gene, **DBI**, is a GABA receptor modulator; its role having been contemplated in schizophrenia [16] the authors concluded that DBI might have a symptom modulatory rather than an etiological role in schizophrenia. Unexpectedly, **MOBP**, myelin-associated oligodendrocyte basic protein and **MBP**, myelin basic protein, both appear in **Table 1B**. While myelin-related abnormalities are one of the hallmarks of schizophrenia, surprisingly, not much functional information is available about MOBP beyond its role in the formation of the myelin sheath [17] and a knockout mouse was phenotypically indistinguishable from the wild type [18]. Nevertheless, the authors argue that MOBP probably has a so far undiscovered function, due to the conservation of several alternatively spliced variants in rat and mouse [18].

Interestingly, several genes in the table are cancer-related. **RRAS,** an oncogene is also involved in neuronal axon guidance; **TMEM219** regulates apoptosis; **CDK2AP1** is a putative oral cancer suppressor; **PSMB10**, a proteasome subunit, is upregulated via the NFKB1 pathway in cancer cells. **S100A1** is also associated with several tumor types and inhibits apoptosis in ventricular cardiomyocytes.

**DGCR6 is** associated with DiGeorge syndrome, a consequence of microdeletions in chromosomal region 22q11.2 and also has increased levels in several tumor lines, including lung and colon adenocarcinomas. **FABP7** plays a role in neurogenesis and is a marker of glioma stem cells [19].

Altogether, 11 of the 20 genes with the greatest numbers of hypermethylated probes for their anti-correlating gene partners (Table 1B) are annotated as schizophrenia-related in Genecards and 10 appear in the 108 loci in [7].

### The connection between the positive and negative hubs

The existence of the two kinds of hub genes with high numbers of positive or negative correlating partners raises the question about their functionality and the underlying neurobiological pathways: are they related or do they form mostly separate networks? To answer this question we constructed three gene networks **(Figure 3)**: two representing the positive and negative correlations only for SYNGR1 and MOBP, respectively (Figure 3a&b), and one (Figure 3c) showing a combined network of the two.

**Figure 3.**
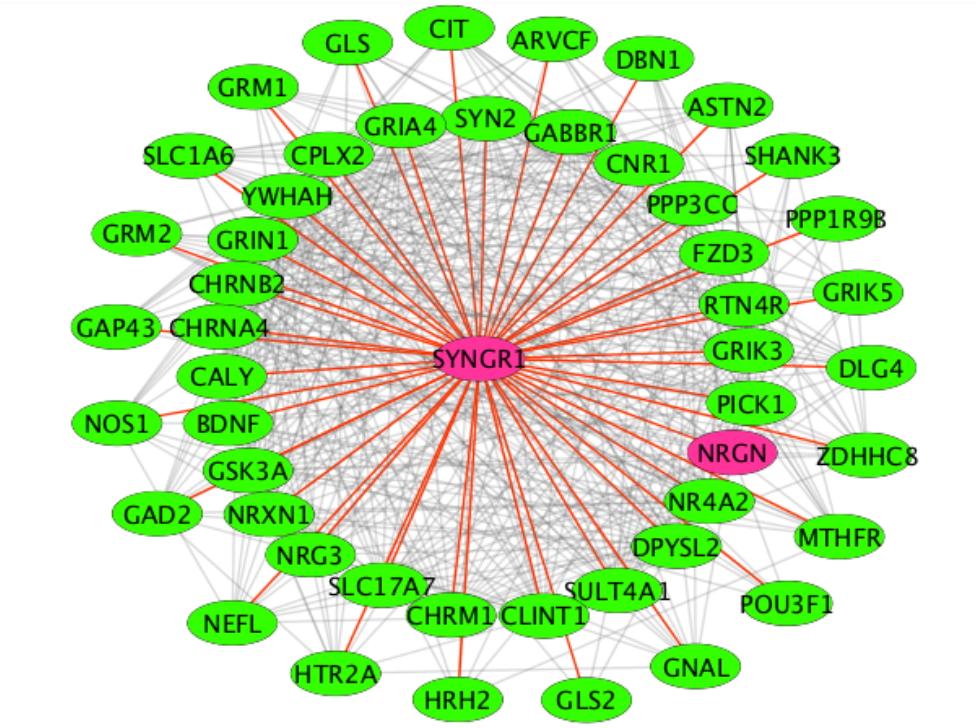
Gene networks of hub genes generated from hypermethylated Malacards-annotated genes. (**A**) SYNGR1-centered, positively correlating gene network. The red-colored edges show correlations with SYNGR1.

**Figure 3 (B).**
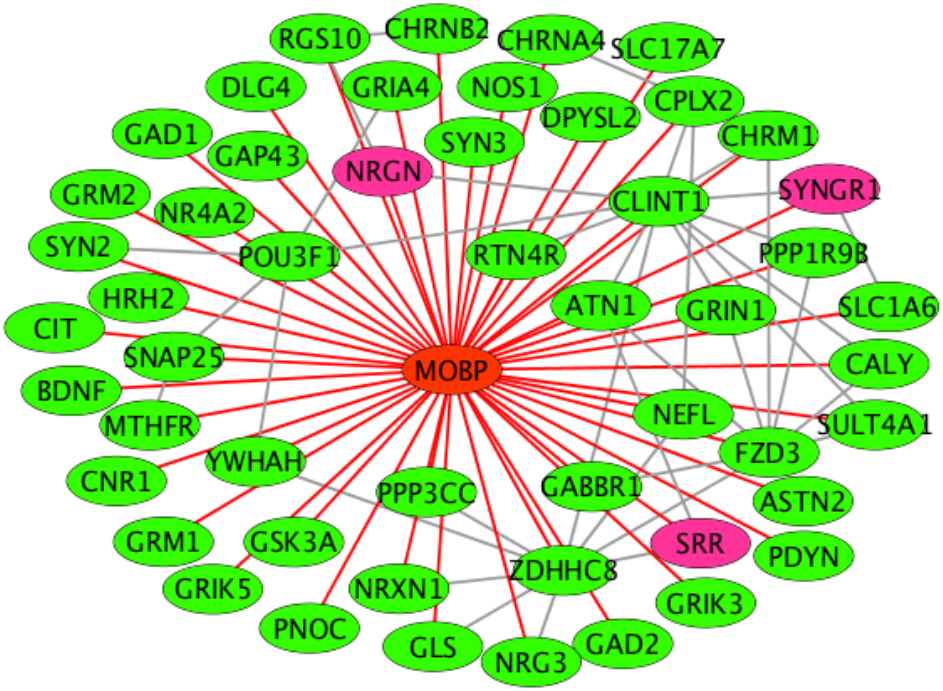
Negatively correlating gene network, centered on MOBP. All the red-colored edges show negative correlations with MOBP. All the other pairwise negative correlations are shown in grey. The purple-colored genes overlap with 108 highly mutated genomic loci in schizophrenics (see text for details).

**Figure 3C.**
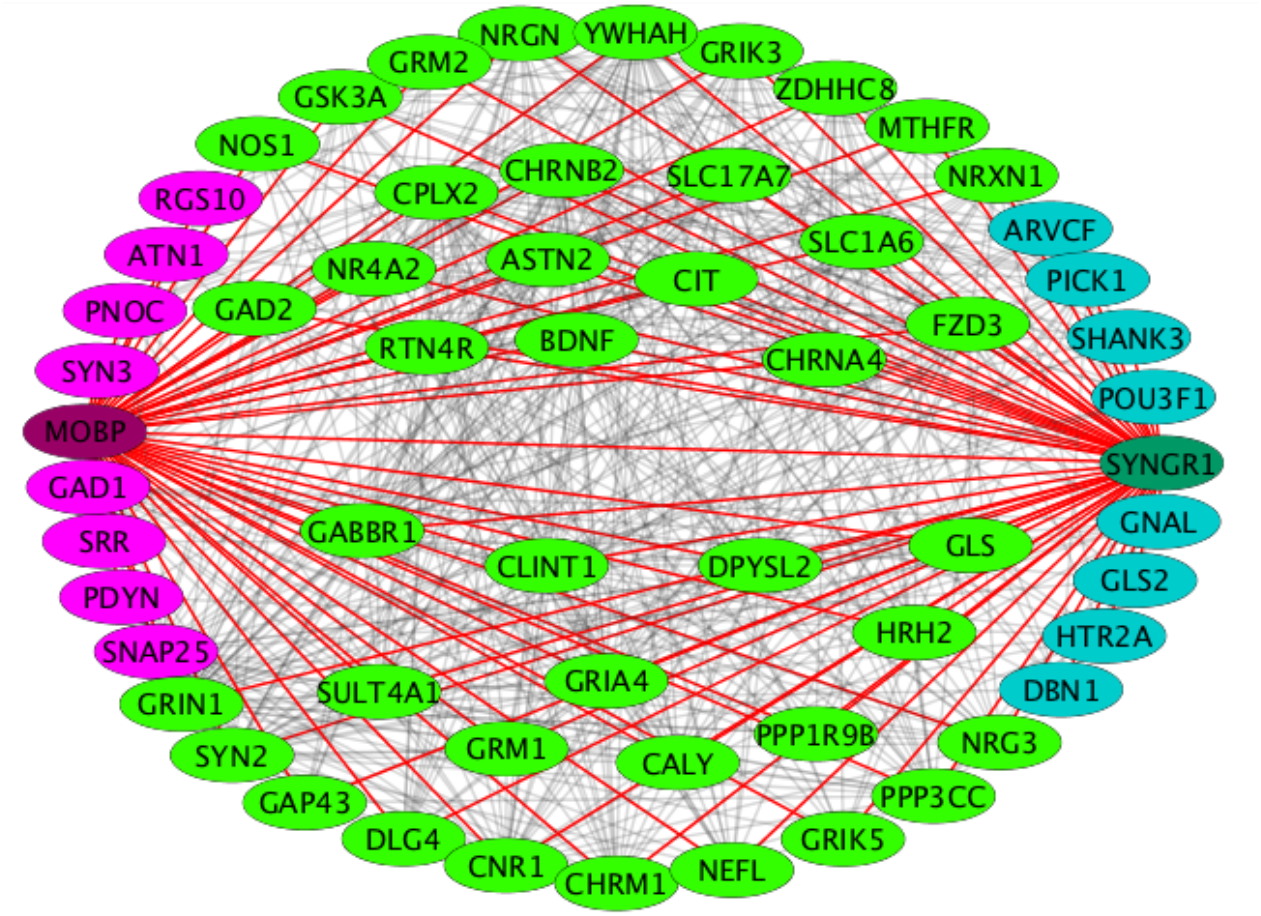
The combined network in **A** and **B**. All light green-colored genes correlate positively with SYNGR1 and negatively with MOBP. The purple-colored and turquoise-colored genes correlate only with MOBP or SYNGR1, respectively.

To reduce the size of the networks we selected only Malacards-annotated schizophrenia-related genes hypermethylated in [8]. SYNGR1 correlates positively with 49 genes (**Figure 3a)**; MOBP also has 49 negatively correlating partners (Figure 3b). The combined network is shown in Figure 3c. Strikingly, 41 genes are shared interacting partners between MOBP and SYNGR1. Each of the two hub genes interacts with only 8 genes not shared with the other hub. Clearly, the positive and negative correlations -and such interactions - form highly interconnected networks that provide synaptic functions, the core functionality of the human brain.

The shared genes reflect remarkably on the nature and most consistently observed features of schizophrenia: 6 shared genes are glutamate receptors (GRIA4, GRIK3, GRIK5, GRIN1, GRM1, GRM2), one GABA receptor (GABBR1), a cannabinoid receptor (CNR1), GAD2, a glutamate decarboxylase, two synapse-related (SYN2, SYT11) and several neuron-specific genes are also present among the shared genes (Figure 3c).

We repeated the selection process replacing the Malacards-genes with the 108-loci genes. This resulted in a similar number of interacting genes that correlate positively with SYNGR1 (50 genes) and negatively with MOBP (48 genes). They share 39 common genes (Figure 4). Biological processes derived from a Gene Ontology analysis of the shared genes either in the Malacards set or the 108-loci set and in both cases shared between SYNGR1 and MOBP and significantly enriched (p-value<0.05) for both sets are shown in Table 2. For both gene sets the top two biological processes with the highest significance are “synaptic transmission” and “modulation of synaptic transmission”.

**Table 2.** Biological processes shared between the Malacards gene set interacting with both SYNGR1 and MOBP in Figure 3C (41 light green-colored genes) and the 108-loci genes interacting with both SYNGR1 and MOBP in Figure 4C (39 light green-colored genes). 12 biological processes in the table are common in the two shared sets, despite the very limited number of actually shared genes (only NRGN is shared besides MOBP and SYNGR1 but these two were not included in the GO analysis).

| GO_identifier | GO_BiologicalProcess | sharedGenes<br>108loci | p_val<br>(108loci) | sharedGenes<br>Malacards | p_val(Malacards<br>Genes) |
| --- | --- | --- | --- | --- | --- |
| GO:0050804 | modulation of synaptic transmission | 7 | 0.00359 | 14 | 1.23e-13 |
| GO:0007268 | synaptic transmission | 9 | 0.00359 | 16 | 8.05e-12 |
| GO:0061564 | axon development | 7 | 0.0318 | 11 | 2.14e-06 |
| GO:0031175 | neuron projection development | 8 | 0.0289 | 11 | 1.08e-05 |
| GO:0048167 | regulation of synaptic plasticity | 4 | 0.0469 | 6 | 4.71e-05 |
| GO:0048666 | neuron development | 9 | 0.0233 | 11 | 4.71e-05 |
| GO:0048812 | neuron projection morphogenesis | 7 | 0.0318 | 9 | 0.000146 |
| GO:0007411 | axon guidance | 6 | 0.0415 | 7 | 0.00128 |
| GO:0030182 | neuron differentiation | 9 | 0.0318 | 9 | 0.00545 |
| GO:0044765 | single-organism transport | 15 | 0.0318 | 15 | 0.0062 |
| GO:0007399 | nervous system development | 14 | 0.0138 | 12 | 0.011 |
| GO:1902578 | single-organism localization | 16 | 0.0289 | 15 | 0.0111 |

**Figure 4 (A).**
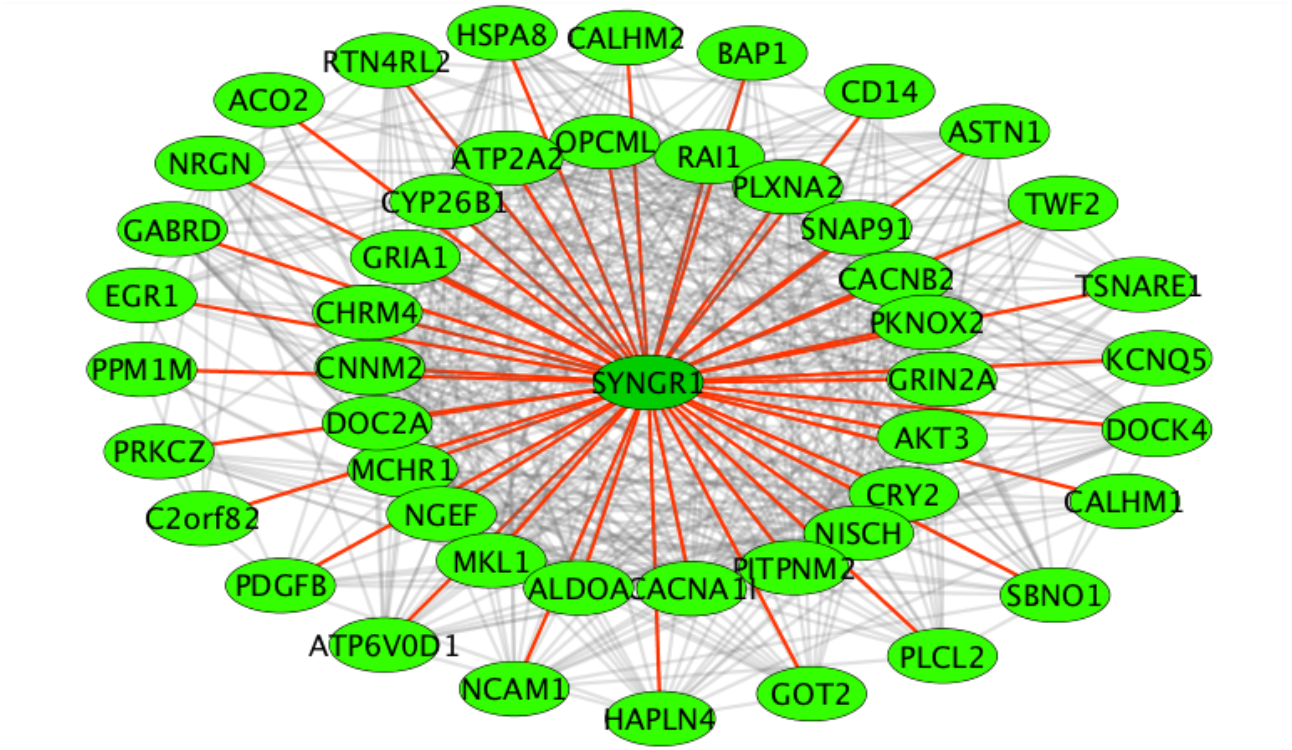
All the positively correlating genes in the 108 loci. The network is again centered on SYNGR1. The red-colored edges show correlations with SYNGR1.

**Figure 4 B.**
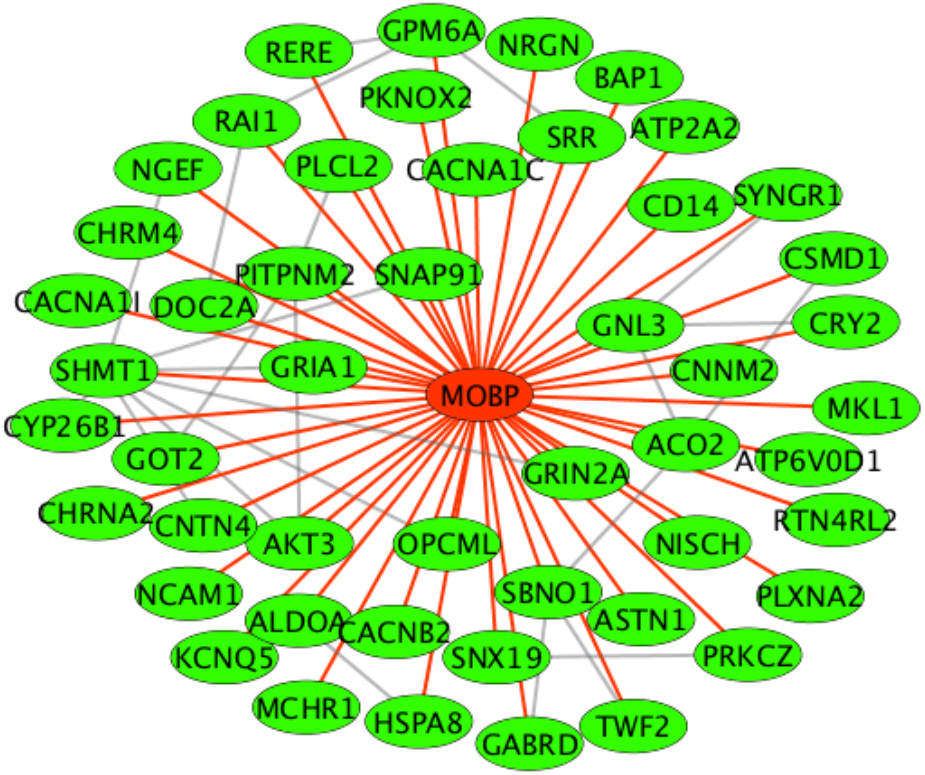
All the genes in the 108 loci that correlate negatively with MOBP (edges colored red) or one another (grey edges).

**Figure 4 C.**
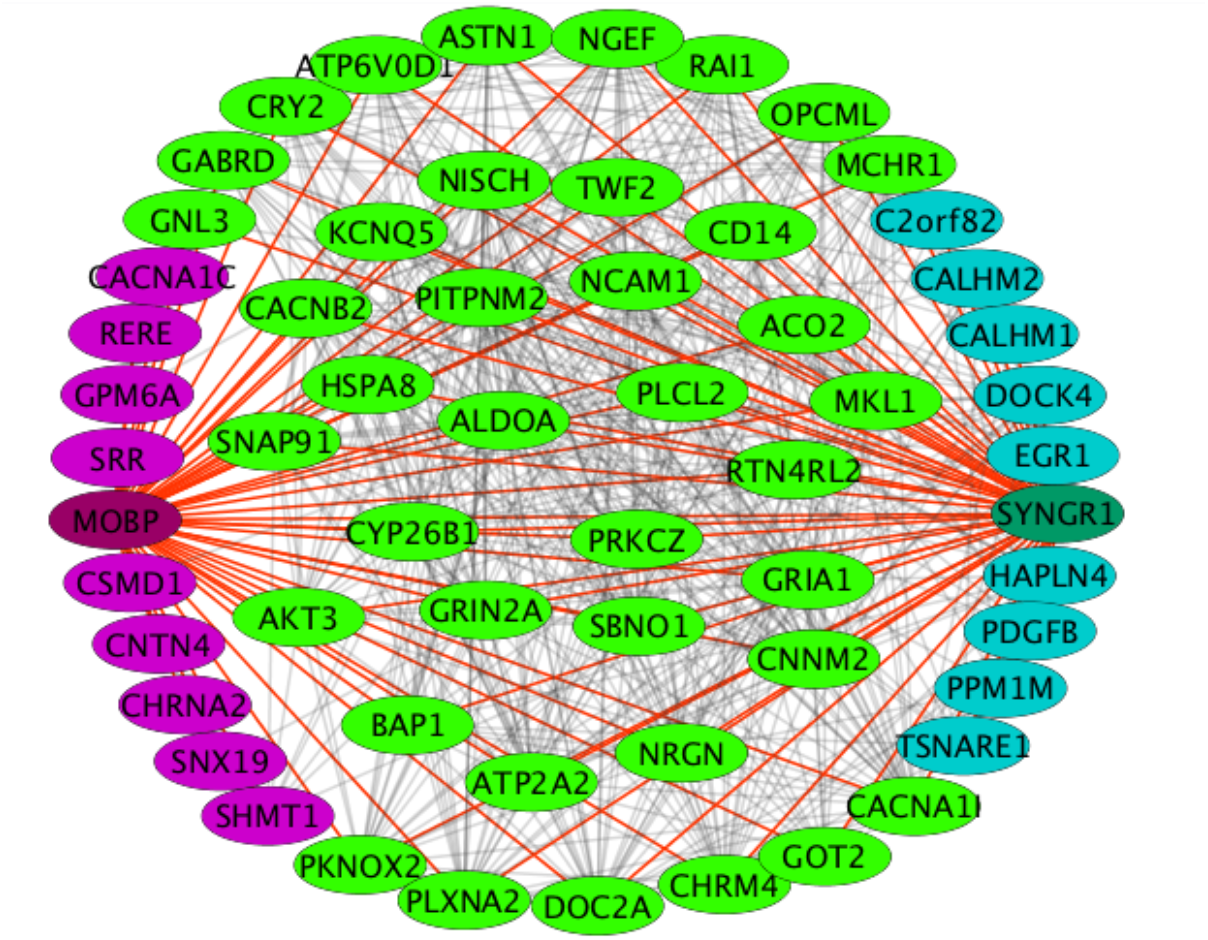
The combined network in **A** and **B**. All light green-colored genes correlate positively with SYNGR1 and negatively with MOBP (edges show in red in both cases). The purple-colored and turquoise-colored genes correlate only with MOBP or SYNGR1 (of the two), respectively.

### Functional analysis of schizophrenia genes correlating with the top 20 hub genes

To get a further insight into the biological processes the top ranking genes and their network neighbors (i.e. their correlating and anti-correlating gene partners) partake in we took the top 20 genes with the most hypermethylated probes for the correlating (and anti-correlating) gene partners and selected those gene neighbors that correlated with *minimum* 19 of the top 20 genes. This resulted in 421 genes for the positively correlating set and 460 genes for the negatively correlating one.

For functional analysis we used an online tool provided by STRING [20] to identify Gene Ontology terms that are over-represented in our gene set. The positive gene set was associated with 278 significantly enriched GO terms for biological processes while the negative set had 151 such GO terms. Interestingly, while the two gene sets share only 117 common genes, i.e. about 1/4^th^ of the total gene number for either set, they also share 117 common biological processes, a much higher fraction for either set (42% of the positive set-derived biological processes and 77% of the negative set-derived processes are shared). The shared GO terms ranked by significance are listed in Table 3. This result also shows that gene networks as a whole are remarkably redundant, i.e. similar functions are often carried out by several functional homologs in the brain. This network view of the genes associated with schizophrenia is also revelatory for the polygenic nature of schizophrenia: once a mutation perturbs the expression of a highly connected hub gene with many interacting partners, this in turn will lead to the perturbation of several hundred or even thousand interacting genes. The differential methylation of several thousands of genes in the prefrontal cortex of schizophrenics might be one such manifestation of the complexity of the disease.

**Table 3.**
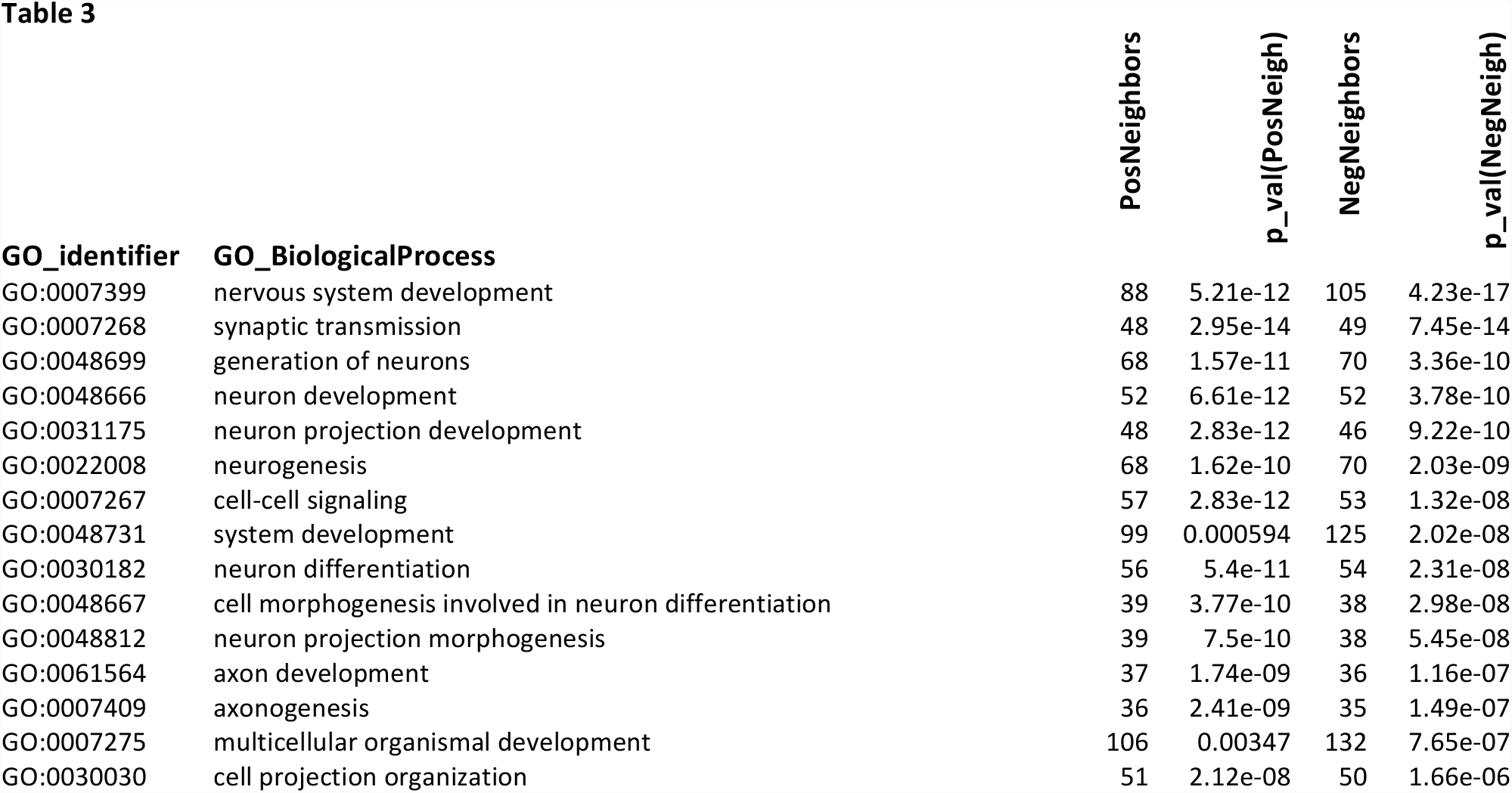
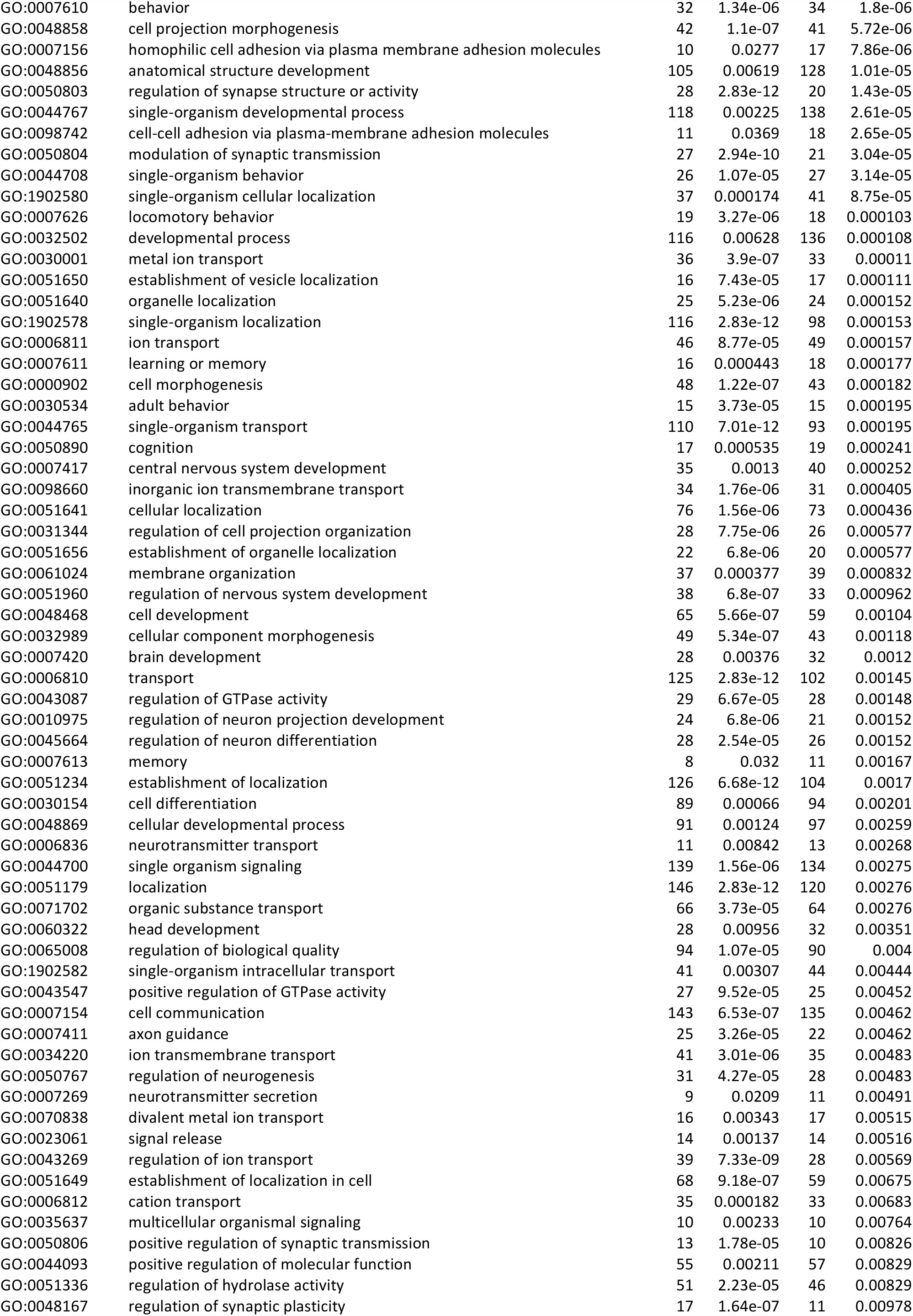
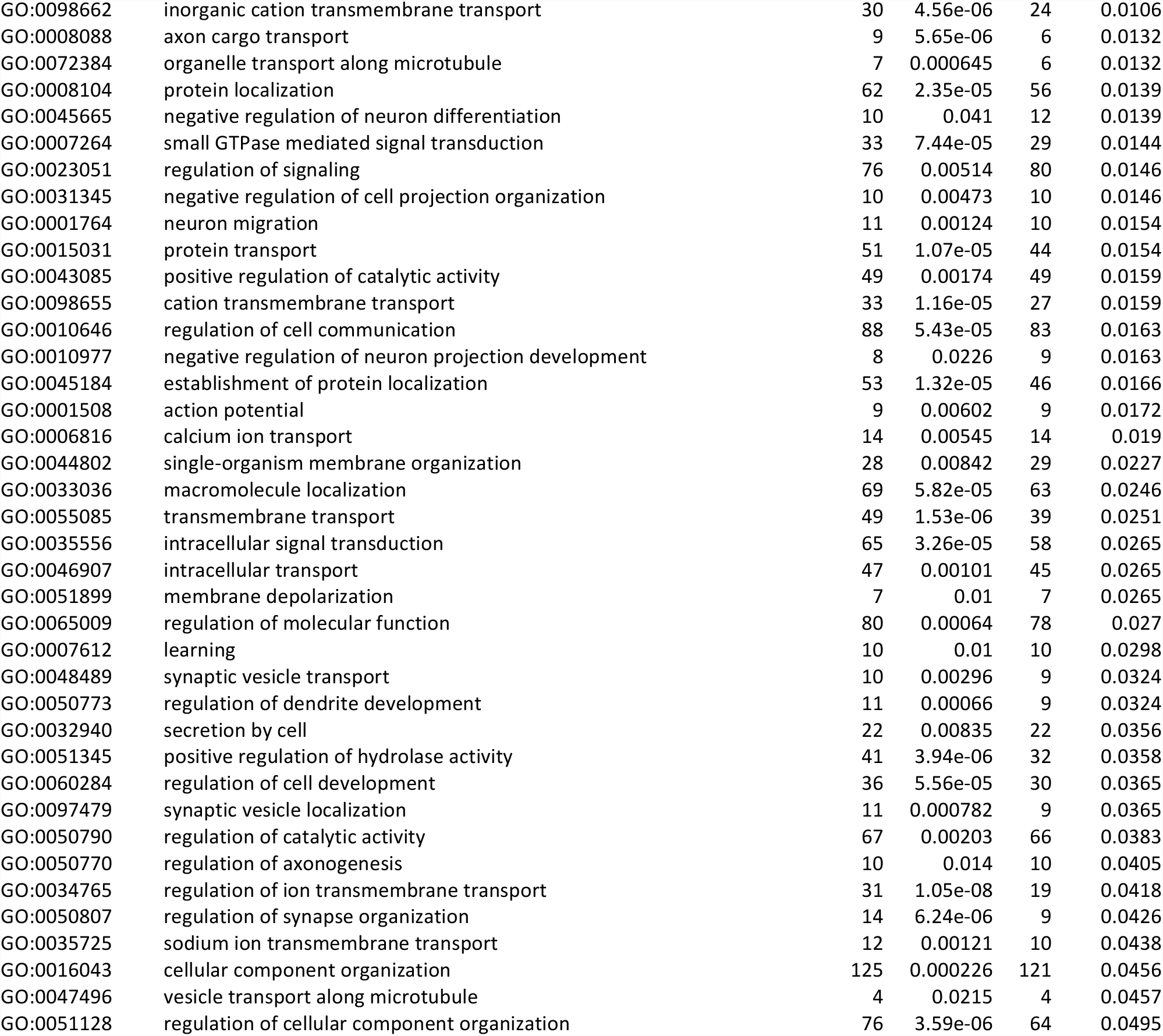
Biological processes shared between the positively correlating and negatively correlating gene neighbors of the top 20 genes in Table 1.

| GO_identifier | GO_BiologicalProcess | PosNeighbors | p_val(PosNeigh) | NegNeighbors | p_val(NegNeigh) |
| --- | --- | --- | --- | --- | --- |
| GO:0007399 | nervous system development | 88 | 5.21e-12 | 105 | 4.23e-17 |
| GO:0007268 | synaptic transmission | 48 | 2.95e-14 | 49 | 7.45e-14 |
| GO:0048699 | generation of neurons | 68 | 1.57e-11 | 70 | 3.36e-10 |
| GO:0048666 | neuron development | 52 | 6.61e-12 | 52 | 3.78e-10 |
| GO:0031175 | neuron projection development | 48 | 2.83e-12 | 46 | 9.22e-10 |
| GO:0022008 | neurogenesis | 68 | 1.62e-10 | 70 | 2.03e-09 |
| GO:0007267 | cell-cell signaling | 57 | 2.83e-12 | 53 | 1.32e-08 |
| GO:0048731 | system development | 99 | 0.000594 | 125 | 2.02e-08 |
| GO:0030182 | neuron differentiation | 56 | 5.4e-11 | 54 | 2.31e-08 |
| GO:0048667 | cell morphogenesis involved in neuron differentiation | 39 | 3.77e-10 | 38 | 2.98e-08 |
| GO:0048812 | neuron projection morphogenesis | 39 | 7.5e-10 | 38 | 5.45e-08 |
| GO:0061564 | axon development | 37 | 1.74e-09 | 36 | 1.16e-07 |
| GO:0007409 | axonogenesis | 36 | 2.41e-09 | 35 | 1.49e-07 |
| GO:0007275 | multicellular organismal development | 106 | 0.00347 | 132 | 7.65e-07 |
| GO:0030030 | cell projection organization | 51 | 2.12e-08 | 50 | 1.66e-06 |
| GO:0007610 | behavior | 32 | 1.34e-06 | 34 | 1.8e-06 |
| GO:0048858 | cell projection morphogenesis | 42 | 1.1e-07 | 41 | 5.72e-06 |
| GO:0007156 | homophilic cell adhesion via plasma membrane adhesion molecules | 10 | 0.0277 | 17 | 7.86e-06 |
| GO:0048856 | anatomical structure development | 105 | 0.00619 | 128 | 1.01e-05 |
| GO:0050803 | regulation of synapse structure or activity | 28 | 2.83e-12 | 20 | 1.43e-05 |
| GO:0044767 | single-organism developmental process | 118 | 0.00225 | 138 | 2.61e-05 |
| GO:0098742 | cell-cell adhesion via plasma-membrane adhesion molecules | 11 | 0.0369 | 18 | 2.65e-05 |
| GO:0050804 | modulation of synaptic transmission | 27 | 2.94e-10 | 21 | 3.04e-05 |
| GO:0044708 | single-organism behavior | 26 | 1.07e-05 | 27 | 3.14e-05 |
| GO:1902580 | single-organism cellular localization | 37 | 0.000174 | 41 | 8.75e-05 |
| GO:0007626 | locomotory behavior | 19 | 3.27e-06 | 18 | 0.000103 |
| GO:0032502 | developmental process | 116 | 0.00628 | 136 | 0.000108 |
| GO:0030001 | metal ion transport | 36 | 3.9e-07 | 33 | 0.00011 |
| GO:0051650 | establishment of vesicle localization | 16 | 7.43e-05 | 17 | 0.000111 |
| GO:0051640 | organelle localization | 25 | 5.23e-06 | 24 | 0.000152 |
| GO:1902578 | single-organism localization | 116 | 2.83e-12 | 98 | 0.000153 |
| GO:0006811 | ion transport | 46 | 8.77e-05 | 49 | 0.000157 |
| GO:0007611 | learning or memory | 16 | 0.000443 | 18 | 0.000177 |
| GO:0000902 | cell morphogenesis | 48 | 1.22e-07 | 43 | 0.000182 |
| GO:0030534 | adult behavior | 15 | 3.73e-05 | 15 | 0.000195 |
| GO:0044765 | single-organism transport | 110 | 7.01e-12 | 93 | 0.000195 |
| GO:0050890 | cognition | 17 | 0.000535 | 19 | 0.000241 |
| GO:0007417 | central nervous system development | 35 | 0.0013 | 40 | 0.000252 |
| GO:0098660 | inorganic ion transmembrane transport | 34 | 1.76e-06 | 31 | 0.000405 |
| GO:0051641 | cellular localization | 76 | 1.56e-06 | 73 | 0.000436 |
| GO:0031344 | regulation of cell projection organization | 28 | 7.75e-06 | 26 | 0.000577 |
| GO:0051656 | establishment of organelle localization | 22 | 6.8e-06 | 20 | 0.000577 |
| GO:0061024 | membrane organization | 37 | 0.000377 | 39 | 0.000832 |
| GO:0051960 | regulation of nervous system development | 38 | 6.8e-07 | 33 | 0.000962 |
| GO:0048468 | cell development | 65 | 5.66e-07 | 59 | 0.00104 |
| GO:0032989 | cellular component morphogenesis | 49 | 5.34e-07 | 43 | 0.00118 |
| GO:0007420 | brain development | 28 | 0.00376 | 32 | 0.0012 |
| GO:0006810 | transport | 125 | 2.83e-12 | 102 | 0.00145 |
| GO:0043087 | regulation of GTPase activity | 29 | 6.67e-05 | 28 | 0.00148 |
| GO:0010975 | regulation of neuron projection development | 24 | 6.8e-06 | 21 | 0.00152 |
| GO:0045664 | regulation of neuron differentiation | 28 | 2.54e-05 | 26 | 0.00152 |
| GO:0007613 | memory | 8 | 0.032 | 11 | 0.00167 |
| GO:0051234 | establishment of localization | 126 | 6.68e-12 | 104 | 0.0017 |
| GO:0030154 | cell differentiation | 89 | 0.00066 | 94 | 0.00201 |
| GO:0048869 | cellular developmental process | 91 | 0.00124 | 97 | 0.00259 |
| GO:0006836 | neurotransmitter transport | 11 | 0.00842 | 13 | 0.00268 |
| GO:0044700 | single organism signaling | 139 | 1.56e-06 | 134 | 0.00275 |
| GO:0051179 | localization | 146 | 2.83e-12 | 120 | 0.00276 |
| GO:0071702 | organic substance transport | 66 | 3.73e-05 | 64 | 0.00276 |
| GO:0060322 | head development | 28 | 0.00956 | 32 | 0.00351 |
| GO:0065008 | regulation of biological quality | 94 | 1.07e-05 | 90 | 0.004 |
| GO:1902582 | single-organism intracellular transport | 41 | 0.00307 | 44 | 0.00444 |
| GO:0043547 | positive regulation of GTPase activity | 27 | 9.52e-05 | 25 | 0.00452 |
| GO:0007154 | cell communication | 143 | 6.53e-07 | 135 | 0.00462 |
| GO:0007411 | axon guidance | 25 | 3.26e-05 | 22 | 0.00462 |
| GO:0034220 | ion transmembrane transport | 41 | 3.01e-06 | 35 | 0.00483 |
| GO:0050767 | regulation of neurogenesis | 31 | 4.27e-05 | 28 | 0.00483 |
| GO:0007269 | neurotransmitter secretion | 9 | 0.0209 | 11 | 0.00491 |
| GO:0070838 | divalent metal ion transport | 16 | 0.00343 | 17 | 0.00515 |
| GO:0023061 | signal release | 14 | 0.00137 | 14 | 0.00516 |
| GO:0043269 | regulation of ion transport | 39 | 7.33e-09 | 28 | 0.00569 |
| GO:0051649 | establishment of localization in cell | 68 | 9.18e-07 | 59 | 0.00675 |
| GO:0006812 | cation transport | 35 | 0.000182 | 33 | 0.00683 |
| GO:0035637 | multicellular organismal signaling | 10 | 0.00233 | 10 | 0.00764 |
| GO:0050806 | positive regulation of synaptic transmission | 13 | 1.78e-05 | 10 | 0.00826 |
| GO:0044093 | positive regulation of molecular function | 55 | 0.00211 | 57 | 0.00829 |
| GO:0051336 | regulation of hydrolase activity | 51 | 2.23e-05 | 46 | 0.00829 |
| GO:0048167 | regulation of synaptic plasticity | 17 | 1.64e-07 | 11 | 0.00978 |
| GO:0098662 | inorganic cation transmembrane transport | 30 | 4.56e-06 | 24 | 0.0106 |
| GO:0008088 | axon cargo transport | 9 | 5.65e-06 | 6 | 0.0132 |
| GO:0072384 | organelle transport along microtubule | 7 | 0.000645 | 6 | 0.0132 |
| GO:0008104 | protein localization | 62 | 2.35e-05 | 56 | 0.0139 |
| GO:0045665 | negative regulation of neuron differentiation | 10 | 0.041 | 12 | 0.0139 |
| GO:0007264 | small GTPase mediated signal transduction | 33 | 7.44e-05 | 29 | 0.0144 |
| GO:0023051 | regulation of signaling | 76 | 0.00514 | 80 | 0.0146 |
| GO:0031345 | negative regulation of cell projection organization | 10 | 0.00473 | 10 | 0.0146 |
| GO:0001764 | neuron migration | 11 | 0.00124 | 10 | 0.0154 |
| GO:0015031 | protein transport | 51 | 1.07e-05 | 44 | 0.0154 |
| GO:0043085 | positive regulation of catalytic activity | 49 | 0.00174 | 49 | 0.0159 |
| GO:0098655 | cation transmembrane transport | 33 | 1.16e-05 | 27 | 0.0159 |
| GO:0010646 | regulation of cell communication | 88 | 5.43e-05 | 83 | 0.0163 |
| GO:0010977 | negative regulation of neuron projection development | 8 | 0.0226 | 9 | 0.0163 |
| GO:0045184 | establishment of protein localization | 53 | 1.32e-05 | 46 | 0.0166 |
| GO:0001508 | action potential | 9 | 0.00602 | 9 | 0.0172 |
| GO:0006816 | calcium ion transport | 14 | 0.00545 | 14 | 0.019 |
| GO:0044802 | single-organism membrane organization | 28 | 0.00842 | 29 | 0.0227 |
| GO:0033036 | macromolecule localization | 69 | 5.82e-05 | 63 | 0.0246 |
| GO:0055085 | transmembrane transport | 49 | 1.53e-06 | 39 | 0.0251 |
| GO:0035556 | intracellular signal transduction | 65 | 3.26e-05 | 58 | 0.0265 |
| GO:0046907 | intracellular transport | 47 | 0.00101 | 45 | 0.0265 |
| GO:0051899 | membrane depolarization | 7 | 0.01 | 7 | 0.0265 |
| GO:0065009 | regulation of molecular function | 80 | 0.00064 | 78 | 0.027 |
| GO:0007612 | learning | 10 | 0.01 | 10 | 0.0298 |
| GO:0048489 | synaptic vesicle transport | 10 | 0.00296 | 9 | 0.0324 |
| GO:0050773 | regulation of dendrite development | 11 | 0.00066 | 9 | 0.0324 |
| GO:0032940 | secretion by cell | 22 | 0.00835 | 22 | 0.0356 |
| GO:0051345 | positive regulation of hydrolase activity | 41 | 3.94e-06 | 32 | 0.0358 |
| GO:0060284 | regulation of cell development | 36 | 5.56e-05 | 30 | 0.0365 |
| GO:0097479 | synaptic vesicle localization | 11 | 0.000782 | 9 | 0.0365 |
| GO:0050790 | regulation of catalytic activity | 67 | 0.00203 | 66 | 0.0383 |
| GO:0050770 | regulation of axonogenesis | 10 | 0.014 | 10 | 0.0405 |
| GO:0034765 | regulation of ion transmembrane transport | 31 | 1.05e-08 | 19 | 0.0418 |
| GO:0050807 | regulation of synapse organization | 14 | 6.24e-06 | 9 | 0.0426 |
| GO:0035725 | sodium ion transmembrane transport | 12 | 0.00121 | 10 | 0.0438 |
| GO:0016043 | cellular component organization | 125 | 0.000226 | 121 | 0.0456 |
| GO:0047496 | vesicle transport along microtubule | 4 | 0.0215 | 4 | 0.0457 |
| GO:0051128 | regulation of cellular component organization | 76 | 3.59e-06 | 64 | 0.0495 |

### Positive hub genes are longer, negative hub genes are shorter than average

We also calculated the protein length and protein disorder statistics for both the top 20 positive and negative hubs. Surprisingly, while the top 20 positive hub proteins were significantly longer (p-value<10^−5^) than the average human protein (894 and 442 amino acids, on average, respectively), the top 20 negative hub proteins were significantly shorter (327 amino acids on average). We did not find their disorder to be significantly different from that of the general human protein population. Likewise, the 108 region-derived proteins among the top 20 positive and negative hubs in Table 1 a & b, were also longer (14 proteins in Table 1a with an average length of 983 amino acids) and shorter (10 proteins in Table 1b with an average length of 261), respectively, than the average human protein (442 aa).

## Discussion

It has been known for at least a decade that myelin has an inhibitory role in axonal regeneration [21] in the CNS. Myelin is dysfunctional in schizophrenia [22] and this dysfunctionality leads to changes in synaptic formation and function, another hallmark of schizophrenia [22]. Several studies have pinpointed genes whose expression is abnormal in schizophrenic brains affecting myelin-related [23] and synapse-related biochemical pathways [24]. However, this is the first time, to our knowledge, that a model for a complete network of gene interactions is presented that would account for both of these recurring anomalies in schizophrenia.

Our gene network has two hubs, one, SYNGR1, with a synaptic function, correlates positively and apparently interacts with a high number of genes that also interact with one another and at the same time interact negatively with MOBP, a myelin gene. Myelin is known to inhibit axonal sprouting, a step considered important for synaptic formation [25]. While neither MOBP, nor MBP are known to have such inhibitory functions, another myelin-related protein, RTN4 (also called Nogo-A), ranking 88th among the negatively correlating genes, does have such a function, inhibiting axon growth [25]. It is tempting to hypothesize that either MOBP or MBP also have such a - so far undiscovered - inhibitory function. MOBP is the 3^rd^ most abundant protein in the CNS myelin and has several alternatively spliced variants. As highlighted by Montague et al. [18] a physiological function for MOBP has not been found yet. Our model with two antagonistic hub genes would also account for the antagonistic relationship between myelin genes and synapse-related functions.

Our model is also supported by the extraordinary enrichment of correlating partners for the genes encoded in the 108 genomic regions with the highest mutation rates in the genomes of schizophrenics identified in [7]. As mentioned above, the median number of correlating partners for those genes whose promoters are located in the 108 loci is 35 times higher for the positively correlating genes than for the rest of the genes in this study and 32 times higher for the negatively correlating genes. The effect size for these comparisons is in the range of 0.8-0.9, making the number of interacting partners, reflecting the centrality of a gene, the single most important factor when considering the biological significance of the individual schizophrenia-related genes and their contribution to the disease. There is a significant but smaller enrichment (2.5 times on average) for those 108-loci genes whose promoters are not, only their enhancers are present in these loci, which raises the intriguing possibility that for most genes in this set the mutations fall into or close to the promoter regions, compromising their functionality as in most cases of schizophrenia the protein sequences are not corrupted by mutations.

The assumption that it is the regulatory regions, not the protein-coding regions that are affected mostly in schizophrenia is also apparent in the fact that despite the strong centrality (“hubness”) of the affected genes we do find surviving phenotypes, which are the patients, exactly. As raised in the introduction, hub genes with a mutation in the coding region would make these mutations lethal in most instances [2].

The robustness of our findings is also supported by the fact that when we replace the Malacards genes with the 108-loci genes in Figure 3 (sharing only SYNGR1, SRR and NRGN as indicated in Figure 3) and analyze the Gene Ontology terms for the 39 genes correlated positively by SYNGR1 and negatively by MOBP, we find that the most significant biological process associated with this gene set is again “synaptic transmission”. This also shows the remarkable redundancy of the gene networks in the human brain. Altogether, beyond providing an intriguing new model for schizophrenia, with more details for the underlining gene networks than before, it is also fascinating and quite fitting that synaptic transmission, perhaps the most complex and dynamic part of the human brain also entails the most genetic complexity, i.e. the most connected gene networks with the highest number of correlating/interacting gene partners.

## Methods

### Determining pairwise gene expression correlations in the human brain

In the first step all pairwise Pearson correlations were determined for those genes expressed in the brain that have expression data in the Allen Brain Atlas (website: brain-map.org) [9, 10]. Gene expression was measured for 50,000 genes in 524 different tissues taken from several compartments of the brains of several individuals spanning the human lifetime between 2 weeks of post-conception and 40 years of age. We used an in-house Perl script to calculate pairwise correlations complemented by correlation data taken from SZDB.org [11]. We filtered the results keeping only pairwise Pearson correlation that were either minimum 0.8 or maximum −0.7. If there were several values for the same gene pair we used the most extreme ones, the highest and lowest values for the positive and negative correlations, respectively.

We mapped all known human genes in 108 highly mutated human genomic regions identified in [7] separately for genes whose promoters or only *cis* regulatory elements fall into these 108 loci. We determined the location of the promoters and cis elements in the human genome using [26].

### Counting correlating gene pairs and their differentially methylated probes

For each gene whose expression correlated (r>=0.8) or anti-correlated (r<=−0.7) with other genes in our data set (16830 and 16762 genes with correlating and anti-correlating partners, respectively) we counted the number of correlating and anti-correlating partners. Using a methylome data set in [8] that recorded 56001 differentially methylated probes between two subgroups of schizophrenia patients we also counted gene partners for each gene that were hypermethylated or hypomethylated (with at least one hypermethylated or hypomethylated probe, respectively) and also the total number of hyper-or hypomethylated probes for the correlating and anti-correlating gene partners.

### Determining the effect size to distinguish between schizophrenia-related and unrelated genes

To determine which measurement distinguishes the best between schizophrenia-related and non-specific genes we calculated the effect size for the 108-loci genes and also for genes annotated by Genecards and Malacards [12, 13] as schizophrenia-related **(Figure 2)** regarding each specific feature mentioned in the previous paragraph (i.e. the number of correlating genes, the number of hypermethylated, hypomethylated correlating genes and the total sum of the differentially methylated probes of the correlating pairs). We repeated the calculations for the negatively correlating pairs as well. To calculate the effect size we used Cliff's delta. Cliff's delta is defined as

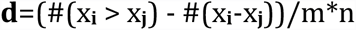

where the two distributions are of size **m** and **n** with items **x_i_** and **x_j_** (with ***i*** running from **1 to m** and ***j*** running from **1 to n**) respectively, and # is defined as the number of times. Cliff's delta shows that out of all possible pairwise comparisons between the numbers in set **A** and set **B** in what proportion will the numbers in set **A** be bigger than in set **B**. Cliff's delta does not require any assumptions about distribution types. All the underlying data are shown in **Supplementary File 2**.

### Functional analysis with the Gene Ontology module of STRING

To carry out functional analysis of the top 20 genes and their network neighbors we used the Gene Ontology (GO) module [14] of the STRING [20] webserver. The server lists all functional categories that are significantly enriched in the provided gene set, and supplies the corresponding p-values. We recorded the biological processes for the top 20 positive and negative genes and also their network neighbors.

### Network visualization, statistical calculations

To visualize the gene correlation networks in Figures 3–4 Cytoscape [27] was used. To carry out t-test calculations and calculate the corresponding p-values we used cpan's Statistics package, a Perl library. Whenever not mentioned explicitly, calculations and data manipulation was carried out with in-house Perl script (available on request from the author).

## Competing interests

None.

## Funding information

This work has been supported by a startup grant at CEITEC.

## Legends to tables

**Supplementary Figure 1.**
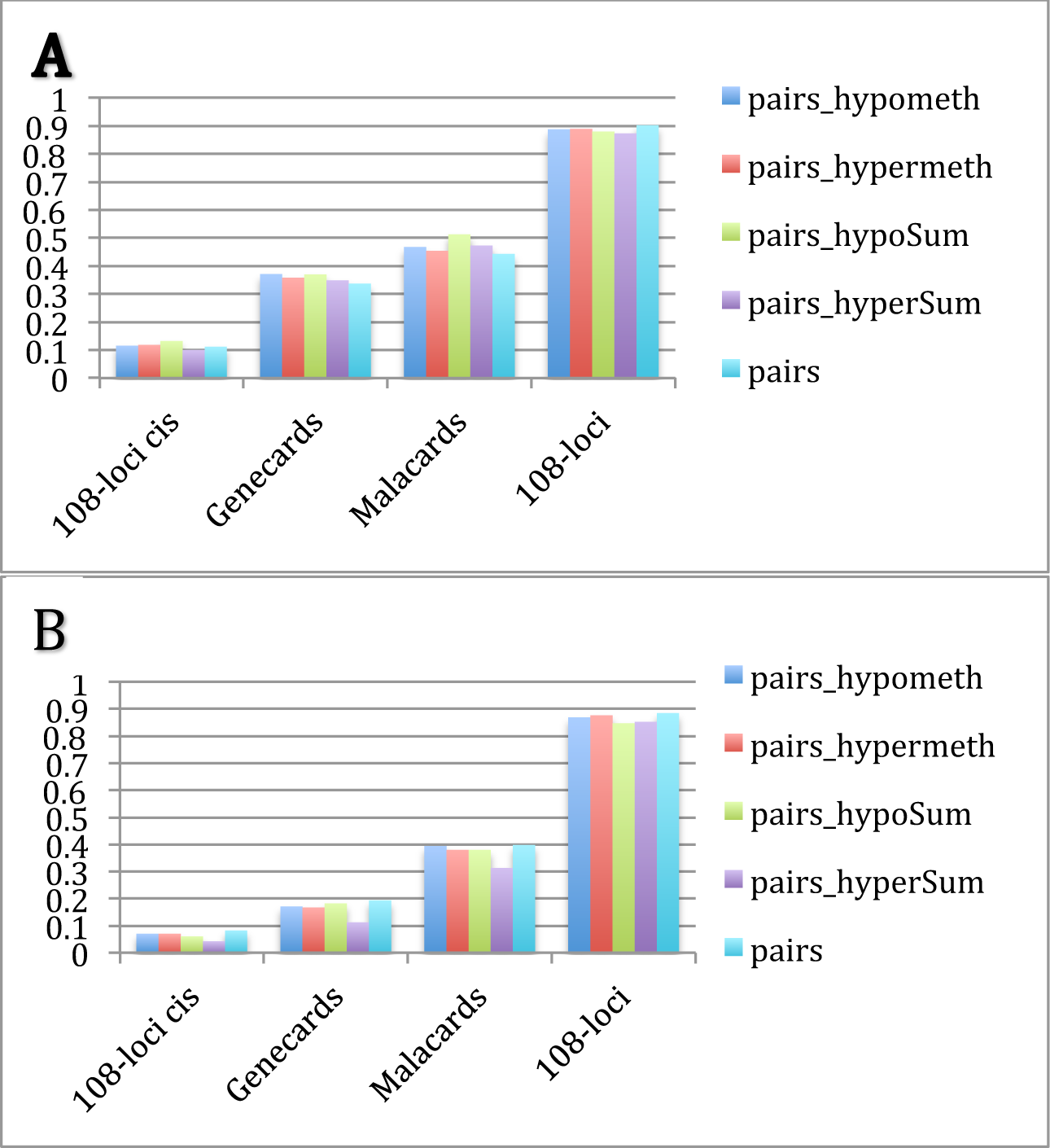
The **effect sizes** for the ratios of gene neighbor numbers between **specific** genes and their **complementary** gene sets (i.e. the rest of the ~16k genes present in the study): 108-loci cis genes; Malacards-annotated schizophrenia genes; Genecards-annotated schizophrenia genes; 108-loci promoter genes. Effect sizes were calculated for 5 different measurements: (i) **pairs**, the total number of correlating gene pairs; (ii) **pairs_hypometh**, the number of correlating gene pairs that are hypomethylated (defined as in at least one probe the gene is hypomethylated); (iii) **pairs_hypermeth**, the number of correlating gene pairs that are hypermethylated; (iv) **pairs_hypoSum**, the total number of hypomethylated probes of the correlating gene pairs; (v) **pairs_hyperSum**, the total number of hypermethylated pairs for the correlating gene pairs. The measurements are the same as in Figure 2 but here they are derived from all pairs in **Supplementary Table 1A & B**, not filtered for pairs of genes that are both differentially methylated in [8]. (**A**) Positively, (**B**) Negatively correlating gene pairs.

**Supplementary table 1.** (A) Gene pairs with a Pearson correlation of minimum 0.8. (B) Gene pairs with a Pearson correlation of maximum −0.7.

**Supplementary table 2.** (A) Summary information for all genes with positively correlating genes in Suppl. table 1A. (B) Summary information for all genes with negatively correlating genes in Suppl. table 1B.

